# Transient transcriptome sequencing: computational pipeline to quantify genome-wide RNA kinetic parameters and transcriptional enhancer activity

**DOI:** 10.1101/659912

**Authors:** Gabriel Villamil, Leonhard Wachutka, Patrick Cramer, Julien Gagneur, Björn Schwalb

## Abstract

In the accompanying chapter (Gressel, Lidschreiber, Cramer), we describe the detailed experimental protocol for transient transcriptome sequencing (TT-seq). TT-seq detects metabolically labeled, newly synthesized RNA fragments genome-wide in living cells. TT-seq can monitor gene activity and the dynamics of enhancer landscapes with great sensitivity, but this requires careful computational analysis of the data. In this manuscript, we present the bioinformatics workflow used to analyze TT-seq data. In particular, we describe pre-processing steps, including a reliable and robust normalization strategy, and several downstream analysis tools that enable the user to quantify RNA synthesis, splicing and degradation activities. Together, these tools form a comprehensive analysis pipeline that can be adapted to almost any TT-seq application.

## 1. Introduction

Compared to standard RNA-seq data, which mainly reveals exonic regions of mRNAs, TT-seq data is much richer, containing many reads in genomic regions that give rise to short-lived, non-coding RNAs such as intronic or enhancer regions. To take advantage of this richness, we have previously developed elaborate tools for the analysis of raw TT-seq data (Gressel et al. 2017; Michel et al. 2017; Schwalb et al. 2016; Zylicz et al. 2019). Computational tools are also available that enable the estimation of RNA synthesis, splicing, and degradation rates, as well as productive transcription initiation rates. Here we summarize and describe the currently available computational tools to analyse TT-seq data. To use these tools, data are required from experiments conducted as described in the accompanying paper (Chapter, Gressel, Lidschreiber, Cramer). Such experiments lead to TT-seq data in the form of short paired-end read sequences of 4sU-labeled, newly synthesized RNA fragments, and equivalent total RNA-seq data derived from the same samples at a similar coverage. To make full use of the data and allow for global normalization, RNA spike-in controls are also necessary. In the following, we provide an overview of data pre-processing steps, the annotation of different RNA transcript classes, and the estimation of transcription- and RNA degradation-related kinetic parameters. We also describe a R-package that combines many of the tools for downstream analysis, including splicing-related kinetic parameters.

## Methods

### I. Pre-processing

#### Read mapping

Begin by demultiplexing the barcoded reads from both TT-seq and RNA-seq experiments and mapping them to the most recent genome reference assembly of your organism. TT-seq read mapping is typically done using STAR (Dobin and Gingeras 2015), with a maximum of two mismatches allowed per 100 bases alignment length. In cases of multi-mapped reads, the single best alignment should be taken. Use Samtools (Li et al. 2009) to quality filter SAM files, whereby alignments with MAPQ smaller than 7 (-q 7) are skipped and only proper pairs (-f 2) are selected. Further data processing may be carried out using the R/Bioconductor environment.

#### Estimating global normalization parameters

We recommend using a RNA spike-in normalization strategy as described (Schwalb et al. 2016) to allow observation of sequencing depth *σ*_*j*_ (global shifts), cross-contamination rate *ϵ*_*j*_ (proportion of unlabeled reads purified in the TT-seq samples), and antisense bias ratio *c*_*j*_ (ratio of spurious reads originating from the opposite strand introduced by the reverse transcription reaction). Read counts (*k*_*ij*_) for spike-ins can be counted using HTSeq (Anders et al. 2015). Global normalization parameters are calculated as follows:

#### Antisense bias ratio *c*_*j*_

Antisense bias ratios are calculated for each sample *j* according to

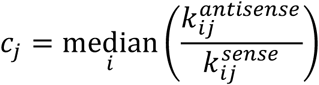

for all available spike-ins *i*.

#### Sequencing depth *σ*_*j*_ and cross-contamination rate *ϵ*_*j*_

Sequencing depths are calculated for each sample *j* according to

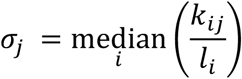

using all available (unlabeled and labeled) spike-ins *i* for the RNA-seq samples and only labeled spike-ins *i* for the TT-seq samples, where *l*_*i*_ defines the length of each spike-in. The cross-contamination rate *ϵ*_*j*_ is calculated for each sample *j* as

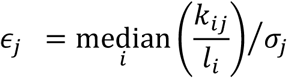

using the unlabeled spike-ins *i* for TT-seq samples. Note that *ϵ*_*j*_ is set to 1 for RNA-seq samples as RNA-seq samples do not undergo 4sU-labeled pull-down purification, and thus have maximal amounts of unlabeled RNA. Note that, sequencing depth *σ*_*j*_ should be standardized by using

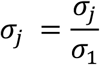

after calculation of the cross-contamination rate *ϵ*_*j*_.

### II. Transcriptome annotation and normalization

#### Definition of transcription units based on the UCSC RefSeq genome assembly (RefSeq-TUs)

For each annotated gene, transcription units are defined as the union of all existing inherent transcript isoforms (UCSC RefSeq). These may be used for general reproducibility assessment and other normalization strategies if applicable.

#### Definition of isoform-independent exonic regions (constitutive exons)

Isoform-independent exonic regions are determined using a model for constitutive exons (Bullard et al. 2010) based on UCSC RefSeq annotation. These can be used to assess the intron enrichment ratio of TT-seq samples which should generally range between 50-90% dependent on the organism or cell type.

#### GenoSTAN annotation of transcription units (TUs)

Annotation of different transcript classes may be done as in (Schwalb et al. 2016). In brief, genome-wide coverage is calculated from all TT-seq fragment midpoints in consecutive 200 bp bins throughout the genome. Binning reduces the number of uncovered positions within expressed transcripts and increases the sensitivity for detection of lowly synthesized transcripts. In order to create a comprehensive annotation, replicate tracks are constructed by taking the maximum of each bin over all corresponding replicates, respectively, regardless of treatment. We train a two-state hidden Markov model with a Poisson-Log-Normal emission distribution to segment the genome into ‘transcribed’ and ‘untranscribed’ states. Consecutive ‘transcribed’ states are joined if its gaps are smaller than 200 bp, are within a validated annotated mRNA or lincRNA, or show uninterrupted coverage supported by all TT-seq samples. Subsequently, TU start and end sites are refined to nucleotide precision by defining borders of abrupt coverage increase or decrease. This is done by finding two consecutive segments in the four 200 bp bins located around the initially assigned start and stop sites via fitting a piecewise constant curve to the TT-seq coverage profiles for all replicates. This can be done for example by using the segmentation method from the R/Bioconductor package “tilingArray” (Huber et al. 2006).

#### GRO-cap or CAGE TSS refinement of TUs (cTUs)

For all TUs *i*, the TU start sites may be refined using GRO-cap or CAGE data by determining the closest non-zero GRO-cap or CAGE signal in a window of 500 bp around the start of the TUs. All TUs without an assigned GRO-cap or CAGE site can either be discarded or used without refined TSS.

#### Transcript sorting

We sort each gene (cTU, TU) into one of the following seven classes: protein-coding (m), long intergenic non-coding (linc), antisense (as), convergent (con), upstream antisense (ua), short intergenic non-coding (sinc), and putative enhancer (e) RNAs. First, (c)TUs reciprocally overlapping by at least 50% with a validated annotated mRNA or lincRNA on the same strand are classified as mRNAs or lincRNAs, respectively. (c)TUs reciprocally overlapping by less than 50% with a validated annotated mRNA or lincRNA on the same strand are not classified. Next, (c)TUs located on the opposite strand of either a mRNA or lincRNA are classified as asRNA if the TSS is located > 1 kb downstream of the sense TSS on the opposite strand, as uaRNA if its TSS is located < 1 kb upstream of the sense TSS, and as conRNA if its TSS is located < 1 kb downstream of the TSS on the opposite strand. Each of the remaining cTUs are classified as sincRNA. Every ncRNA (sincRNA, asRNA, conRNA or uaRNA) can subsequently be re-classified as eRNA if its TSS falls into an enhancer state based on genome state classification data, such as (Zacher et al. 2017), or if its TSS exhibits a high (>1) ratio of H3K4me1 to H3K4me3.

#### TT-seq and RNA-seq data processing with global normalization parameters

Read counts or coverage *k*_*ij*_ for all features ((c)TUs, constitutive exons or RefSeq-TUs) are normalized and corrected using the parameters calculated as described above for antisense bias *c*_*j*_, sequencing depth *σ*_*j*_ and cross-contamination rate *ϵ*_*j*_ as follows.

#### Antisense bias correction

The real number of read counts or coverage *s*_*ij*_ for transcribed unit *i* in sample *j* is calculated as

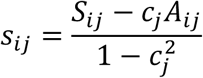

where *S*_*ij*_ and *A*_*ij*_ are the observed numbers of read counts or coverage on the sense and antisense strand, respectively.

#### Sequencing depth normalization and cross-contamination correction

The antisense bias corrected read counts or coverage *s*_*ij*_ of transcribed unit *i* in sample *j* is normalized for sequencing depth and corrected for cross-contamination as

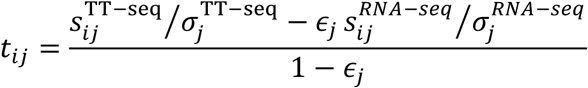

Note that, *σ*_*j*_ can be balanced between replicates via classical size factor normalization (Anders and Huber 2010) to gain statistical power in the differential expression analysis.

#### Read counts per kilobase (RPK)

RPKs are calculated upon antisense bias corrected read counts *k*_*ij*_ falling into the region of a (c)TU, constitutive exon or RefSeq-TU divided by its length in kilobases.

#### Calculation of the number of transcribed bases

The number of transcribed bases *tb*_*ij*_ for all samples are calculated as the sum of the coverage of evident (sequenced) fragment parts (read pairs only) for all fragments in addition to the sum of the coverage of non-evident fragment parts for fragments with an inner mate interval not entirely overlapping an annotated intron. The number of transcribed bases *tb*_*ij*_ can later be used to calculate estimates of the productive initiation frequency. Coverages are calculated upon antisense bias corrected number of transcribed bases *tb*_*ij*_ falling into the region of a (c)TU or RefSeq-TU divided by its length in bases.

### III. Estimation of kinetic parameters

#### Molecular weight conversions

The known sequence and mixture of the utilized spike-ins allows to calculate a conversion factor to RNA amount per cell [cell^-1^] given their molecular weight assuming perfect RNA extraction. The number of spike-in molecules per cell *N* [cell^-1^] is calculated as

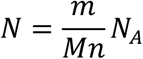

with the mass *m* [g] per spike-in, the number of cells *n*, the Avogadro constant *N*_*A*_ 6.02214085774·10^23^ [mol^-1^] and molar-mass (molecular weight) of the spike-ins *M* [g mol^-1^] calculated as

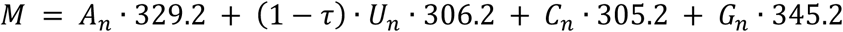

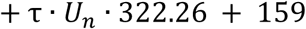

where A_n_, U_n_, C_n_, G_n_ are the number of each respective nucleotide within each spike-in polynucleotide. τ is set to 0.1 in case of a labeled spike-in and 0 otherwise (τ · *U*_2_ corresponds to the number of 4sU nucleotides, 4*sU*_2_). The addition of 159 to the molecular weight takes into account the molecular weight of a 5’ triphosphate. Provided the above the conversion factor to RNA amount per cell [cell^-1^] can be calculated as

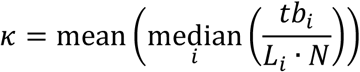

for all labeled spike-in species *i* with length *L*_*i*_.

#### Estimation of productive initiation frequency *I*

For each (c)TU or RefSeq-TU *i* the productive initiation frequency *I*_*i*_ [cell-1min-1] is calculated using the antisense bias corrected number of transcribed bases *tb*_*ij*_ as

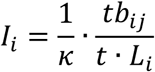

with labeling duration *t* [min] and length *L*_*i*_. For RefSeq-TUs, *tb*_*ij*_ and *L*_*i*_ may be restricted to regions of non-first constitutive exons. This is to ensure to not be biased by alternative splicing.

#### Estimation of RNA synthesis rates *μ*_*ij*_ and degradation rates *λ*_*ij*_

Read counts per base are calculated upon antisense bias corrected read counts *s*_*ij*_ for gene *i* in sample *j* (TT-seq and RNA-seq samples) falling into the region of a feature ((c)TUs, constitutive exons or RefSeq-TUs) divided by its sequencing depth *σ*_*j*_ and length *l*_*i*_ as

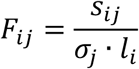

Fraction-specific read counts *F*_*ij*_ are from here on referred to as *L*_*ij*_ for TT-seq or *T*_*ij*_ for RNA-seq samples. Synthesis rates *μ*_*ij*_ and degradation rates *λ*_*ij*_ are subsequently calculated assuming first-order kinetics as in (Miller et al. 2011) using the following equations:

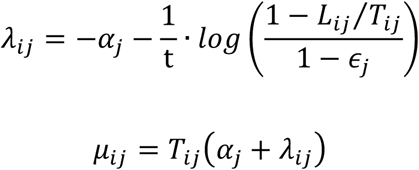

or

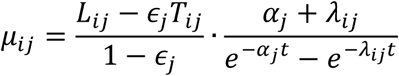

where t is the labeling duration in minutes and *α* is the growth rate (dilution rate, i.e. the reduction of concentration due to the increase of cell volume during growth) defined as

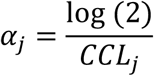

with cell cycle length *CCL*_*j*_ [min].

#### Calculation of calibrated RNA synthesis rates 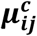

For each TU *i* the antisense bias corrected number of transcribed bases 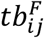 may be used to calculate the calibrated synthesis rate 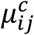 [cell^-1^min^-1^] as

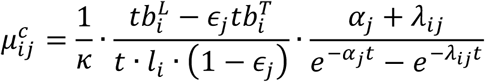

where t is the labeling duration in minutes and *l*_*i*_ the length of the respective feature. This equation utilizes the added spike-ins to adjust synthesis rates to units of nucleotide bond formation per cell per minute.

### IV. Estimation of RNA synthesis and degradation rates using the R package rCube

#### Overview

The R package rCube (**R**NA metabolism **R**ates in **R**) provides functionalities for the estimation of RNA metabolism rates from TT-seq data. Specifically, TT-seq and total RNA-seq data are used to model RNA synthesis, splicing, and degradation rates based on first-order kinetics. Functionalities are provided to extract read counts from BAM files for various genomic features of interest, including splice junctions, and constitutive exons. Moreover, the rCube package implements various sample normalization methods.

#### Installation and loading of the package

The rCube package is installed from the Github repository via the following R commands:

~~~
library(‘devtools’)
install_github(‘gagneurlab/rCube’)
~~~

Before starting, the package must be loaded by:

~~~
library(‘rCube’)
~~~

#### Defining genomic features

The package supports the estimation of kinetic parameters by first-order kinetics for various genomic features. On the one hand, kinetic parameters can be fitted for entire genes based on constitutive exons (Michel et al. 2017). To load the coordinates of these exons and their grouping by genes, an object of the class GRanges (from the Bioconductor GenomicRanges package) is created by parsing a file in GTF format containing the gene annotations. This is achieved by calling the `createConstitutiveFeaturesGRangesFromGRanges` function as follows:

~~~
*library(rtracklayer)*
*granges <-import(‘path to gtf file’)*
constitutiveExons <-
createConstitutiveFeaturesGRangesFromGRanges(*granges*)
~~~

The package rCube also supports the fitting of kinetic parameters for individual nucleotide bonds relevant for splicing kinetics: donor sites, acceptor sites, and exon junction bonds (Wachutka and Gagneur 2017). To this end, a GRanges object with all intron coordinates needs to be provided. This can be done by parsing a file in GTF format containing only introns as follows:

~~~
*library(rtracklayer)*
*granges <-import(‘path to gtf file of introns’)*
~~~

Alternatively, introns can be *de novo* identified using the split reads from RNA-seq data. This is achieved by:

~~~
#bamfiles is a vector of BAM filenames
*granges <-createJunctionGRangesFromBam(bamFiles)*
~~~

#### The rCubeExperiment class

The main class used by rCube is called rCubeExperiment. It consists of:

- a genomic feature annotation of class GRanges as produced in the preceding code,
- a sample information table of class data.frame, and - an assay or matrix containing read counts for all annotated regions in the corresponding samples.

Objects of the class rCubeExperiment are used as input for the whole workflow, including read counting, normalization, dispersion and rate estimation. Most of these steps return an updated and extended rCubeExperiment object. Experimental sample information in the colData of the rCubeExperiment object should contain the following columns:

- sample: A unique sample name
- LT: A factor, stating whether the sample was labeled (‘L’ for TT-seq) or total RNA (‘T’ for total RNA-seq)
- condition: A factor which distinguishes different experimental conditions (but not the sequencing type, e.g. genetic background, stimulation, etc.)
- labelingTime: A numeric value indicating the labeling time for each labeled sample
- replicate: A factor giving replicate information (if no replicates were used, use the same value for all samples)
- filename: An optional string specifying the filename of the corresponding sample

This information can be either provided by a manually set up data.frame. Alternatively, this information can be extracted from the BAM-file names, if they follow the following naming convention: {condition}_{L|T}_{labelingTime}_{replicate}.bam

~~~
#bamfiles has to be a vector of BAM filenames
rCubeCounts **<-** setupExperiment(granges, files**=**bamFiles)
~~~

#### Counting reads

After setting up the experiment, reads mapping to the different genomic features (e.g. all constitutive exons grouped by gene) are obtained by calling

~~~
rCubeCounts <-countFeatures(rCubeCounts)
~~~

Similarly, counts of reads mapping to the donor site, acceptor site or exon-exon bond of an intron by calling:

~~~
rCubeCounts <-countJunctions(rCubeCounts)
~~~

#### Sample normalization and cross-contamination

Sample normalization and estimation of the amount cross-contamination are estimated using labeled and unlabeled spike-ins. To count the spike-in reads, rCube relies on an artificial file in the GFT format containing the spike-in coordinates and specifying their corresponding labeling state:

~~~
#bamfiles has to be a vector of BAM filenames
spikeinsGtf **<-** import(‘spikeins.gtf’)
rCubeSpikein **<-** setupExperimentSpikeins(
             spikeinsGtf,
             files **=** bamFiles,
             labelingState **=** c(‘L’,’U’,’L’,’U’,’L’,’U’)
)
rCubeSpikein **<-** countSpikeins(rCubeSpikein)
~~~

The package rCube offers three different normalization schemes that can be chosen in the `estimateSizeFactors` functions by the parameter method, namely: ‘spikeinGLM’, ‘spikeinMean’, and ‘spikeinMedian’. The ‘spikeinMean’ and ‘spikeinMedian’ normalization methods estimate size factors based on the read distributions of labeled spike-ins among the different samples. The spikeinGLM method is based on generalized linear model fit to the spike-in data (Michel et al. 2017). In general, all three methods give highly correlated size factors. Given the spike-in read counts one can easily use them to derive the necessary sample normalization factors by a call to the function `estimateSizeFactors`:

~~~
rCubeCounts <-estimateSizeFactors(rCubeCounts, rCubeSpikein,
method=“spikeinGLM”)
~~~

#### Estimating the dispersion

Read counts in RNA-seq samples show fluctuations due to biological or technical variations which lead to noise larger than the Poisson noise (over-dispersion (Anders and Huber 2010)). To take the over-dispersion into account, read counts in rCube are modelled using the negative binomial distribution. The wrapper function estimateSizeDispersions offers different methods to estimate negative binomial distribution dispersion parameters. To more accurately estimate the dispersion, we advise to perform replicates for each labeling duration and condition. The call to the `estimateSizeDispersions` function reads:

~~~
rCubeCounts <-estimateSizeDispersions(rCubeCounts,
method=“Replicate”)
~~~

In absence of replicates, the `estimateSizeDispersions` function can be called with the parameter method set to ‘DESeqDispGeneEst’, which estimates dispersion using the read count variations across the whole dataset.

#### Fitting rates

Next, RNA synthesis and degradation rate for each feature and condition can be fitted. The rCube package supports two different experimental designs.

a. The experiment consists of a labeling time series, i.e. a set of TT-seq samples with different labeling times and complementary total RNA-seq samples. Here, a first order kinetic model is fitted to the time series (Eser et al. 2016).
b. The experiment consists of total RNA-seq data and TT-seq data only for one short labeling time. In this case, synthesis and degradation rates can be estimated assuming that no significant degradation has occurred during the short labeling time (Michel et al. 2017; Schwalb et al. 2016).

Depending on the experimental design one can use the method ‘series’ for experiments of type **(A)** or ‘single’ for experiments of type **(B)**. One can further obtain replicate-specific rate estimates, for instance to investigate the reproducibility of the results. The following code estimates results for a type **(A)** experiment for replicate 1 and 2 individually and also results for a joint estimation based on both replicates.

~~~
rates <-estimateRateByFirstOrderKinetics(rCubeCounts,
replicate=c(1,2,1:2), method=“series”)
~~~

After the fitting is done the results can be extracted from the result object using the following accessor functions:

~~~
# feature information
rowRanges(rates)
# sample information
colData(rates)
# rates
assay(rates)
~~~

## Acknowledgements

We thank Carina Demel (Cramer laboratory) for help with the R package rCube. PC was funded by Advanced Grant TRANSREGULON of the European Research Council and the Volkswagen Foundation. This manuscript will be published as a chapter in Methods in Molecular Biology.

## Figure

**Figure 1.**
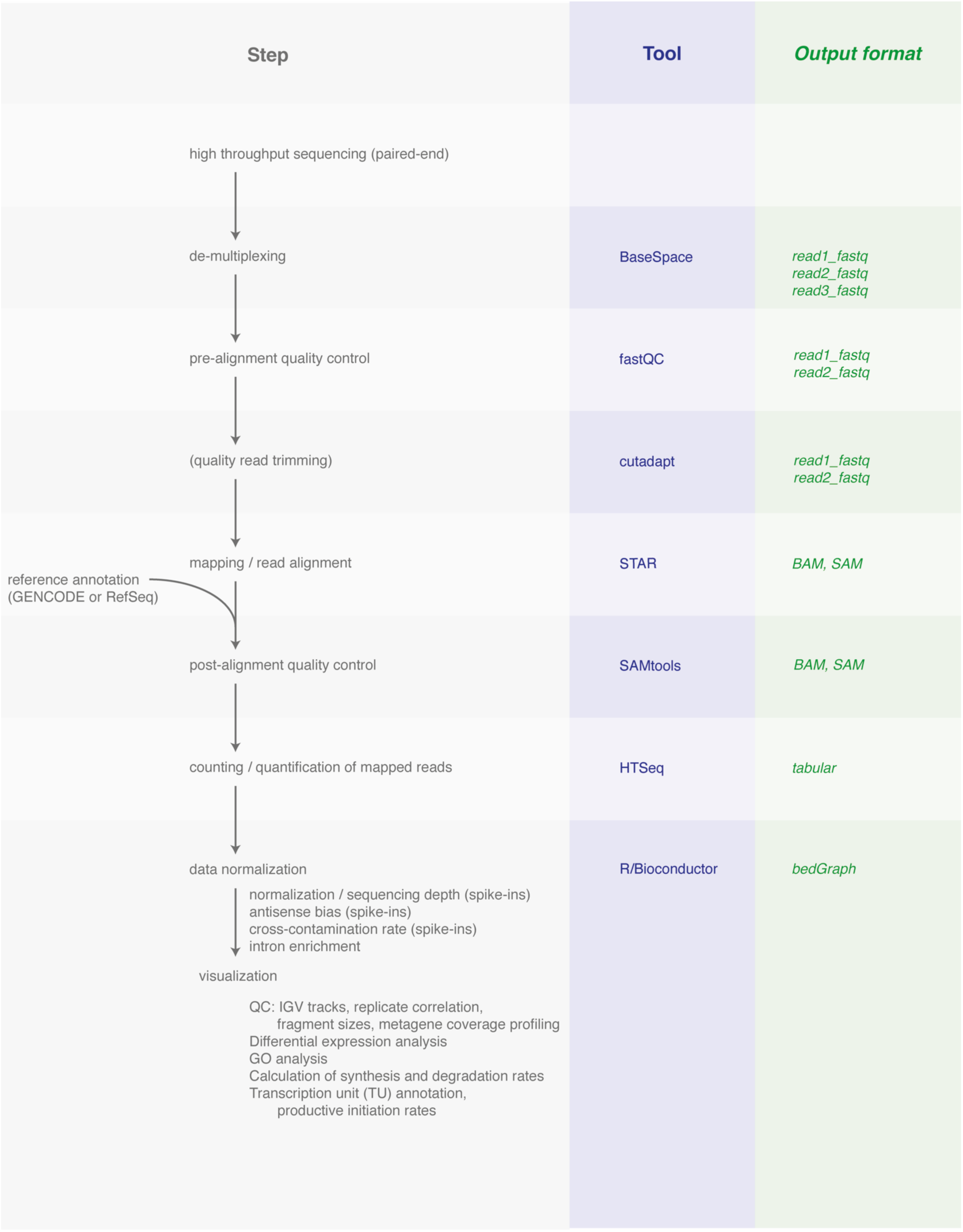
Schematic overview of computational strategy for transient transcriptome sequencing (TT-seq) to measure short lived newly synthesized RNA. For details refer to main text.

## References

Anders S, Huber W (2010) Differential expression analysis for sequence count data Genome Biol 11:R106 doi:10.1186/gb-2010-11-10-r106

Anders S, Pyl PT, Huber W (2015) HTSeq--a Python framework to work with highthroughput sequencing data Bioinformatics 31:166–169 doi:10.1093/bioinformatics/btu638

Bullard JH, Purdom E, Hansen KD, Dudoit S (2010) Evaluation of statistical methods for normalization and differential expression in mRNA-Seq experiments BMC Bioinformatics 11:94 doi:10.1186/1471-2105-11-94

Dobin A, Gingeras TR (2015) Mapping RNA-seq Reads with STAR Curr Protoc Bioinformatics 51:11 14 11–19 doi:10.1002/0471250953.bi1114s51

Eser P et al. (2016) Determinants of RNA metabolism in the Schizosaccharomyces pombe genome Mol Syst Biol 12:857 doi:10.15252/msb.20156526

Gressel S, Schwalb B, Decker TM, Qin W, Leonhardt H, Eick D, Cramer P (2017) CDK9-dependent RNA polymerase II pausing controls transcription initiation Elife 6 doi:10.7554/eLife.29736

Huber W, Toedling J, Steinmetz LM (2006) Transcript mapping with high-density oligonucleotide tiling arrays Bioinformatics 22:1963–1970 doi:10.1093/bioinformatics/btl289

Li H et al. (2009) The Sequence Alignment/Map format and SAMtools Bioinformatics 25:2078–2079 doi:10.1093/bioinformatics/btp352

Michel M et al. (2017) TT-seq captures enhancer landscapes immediately after T-cell stimulation Mol Syst Biol 13:920 doi:10.15252/msb.20167507

Miller C et al. (2011) Dynamic transcriptome analysis measures rates of mRNA synthesis and decay in yeast Mol Syst Biol 7:458 doi:10.1038/msb.2010.112

Schwalb B et al. (2016) TT-seq maps the human transient transcriptome Science 352:1225–1228 doi:10.1126/science.aad9841

Wachutka L, Gagneur J (2017) Measures of RNA metabolism rates: Toward a definition at the level of single bonds Transcription 8:75–80 doi:10.1080/21541264.2016.1257972

Zacher B, Michel M, Schwalb B, Cramer P, Tresch A, Gagneur J (2017) Accurate Promoter and Enhancer Identification in 127 ENCODE and Roadmap Epigenomics Cell Types and Tissues by GenoSTAN PLOS One 12:e0169249 doi:10.1371/journal.pone.0169249

Zylicz JJ et al. (2019) The Implication of Early Chromatin Changes in X Chromosome Inactivation Cell 176:182–197 e123 doi:10.1016/j.cell.2018.11.041

